# Cell explosions: single cell death triggers an avoidance response in local populations of the ecologically prominent phytoplankton genus *Micromonas*

**DOI:** 10.1101/740605

**Authors:** Richard Henshaw, Jonathan Roberts, Marco Polin

## Abstract

The global phytoplankton community, comprised of aquatic photosynthetic organisms, is acknowledged for being responsible for half of the global oxygen production Prominent among these is the pico-eukaryote *Micromonas commoda* (formally *Micromonas pusilla* of the genus *Micromonas*), which can be found in marine and coastal environments across the globe. Cell death of phytoplankton has been identified as contributing to the largest carbon transfers on the planet moving 10^9^ tonnes of carbon in the oceans every day. During a cell death organic matter is released into the local environment which can act as both a food source and a warning signal for nearby organisms. Here we present a novel motility response to single cell death in populations of *Micromonas sp*., where the death of a single cell releases a chemical patch triggers surrounding cells to escape the immediate affected area. These so-called “burst events” are then modelled and compared with a spherically symmetric diffusing patch which is found to faithfully reproduce the observed behaviour. Finally, laser ablation of single cells reproduces the observed avoidance response, confirming that *Micromonas sp*. has evolved a specific motility response in order to escape harmful environments for example nearby predator-prey interactions or virus lysis induced cell death.

## INTRODUCTION

There are over 100,000 recorded algal species in a wide variety of environments such as soil, freshwater and oceans. These species cover a broad spectrum of size magnitudes from the microscopic *Ostrecoccus tauri* [1] (≤ 0.8*μ*m) up to giant 50m *Macrocystis pyrifera*. One reason interest in these organisms is continually growing is due to the myriad of biotechnical applications such as: biofuel production, pollution indicators, hydrogen production and cosmetics [2–6]. Algae are also commercially attractive as they can be cultivated on industrial scales in almost any environment and so provides an almost unparalleled conversion rate of solar energy to carbon rich molecules whilst exerting and almost negligible pressure on arable land.

Despite many algal organisms being on the scale of microns, they have a profound influence that extends far beyond their immediate surroundings to encompass the entire global ecosystem. Many algae species form part of the phytoplankton community – a collection of aquatic photosynthetic organisms which are also estimated for being responsible for over half the global oxygen production. These organisms also affect the global carbon cycle since approximately half the net primary productivity of the entire biosphere is credited to the oceans, currently estimated at 50 Pg.C.yr^-1^ or approximately 50 billion metric tonnes of carbon every year. Living organisms in the oceans contribute to a total carbon mass of 1 —2 Pg.C, a quarter of which is attributed to phytoplankton [7, 8]. This pales in contrast to the mass of the non-living carbon in the oceans which is currently estimated as at least 1,000 Pg.C, the majority of which is defined as “dissolved organic matter” (DOM) [9]. The amount of dissolved organic carbon (DOC) is currently estimated to be in the same range of the quantity of carbon in the atmosphere [10].

This organic material is not only a significant portion of the global carbon cycle but is also a food source for many marine microorganisms [11, 12], party due to how DOC does not sediment (unlike particulate carbon) and so remains available at differing heights in the water column. Phytoplankton act as a source for DOM in the oceans when they release material into the oceans [13–16], for example from extracellular release due to photosynthesis or when a cell dies which provides trace metals and other organic nutrients and forms a crucial part of nutrient recycling [17–19]. Cell death can be triggered through several means such as through viral lysis [20–22] and other accidental means i.e. rapid changes in the local pH or severe local mechanical stresses. Some unicellular eukaryotes are even capable of regulated cell death (RCD) [21] or programmed cell death (PCD) [23] where the organism has developed dedicated molecular machinery which can be triggered by non-environmental conditions (i.e. to promote tissue turnover) or in response to some external stimuli. In this case, some organisms are able to release molecules (called damage-associated molecular patterns, or DAMPs for short) to alert other nearby cells to this potential threat e.g. a predator feeding or a local viral lysis.

One of the most prominent phytoplankton is the pico-eukaryote *Micromonas* (a genus originally containing a single species *Micromonas pusilla* until a recent reclassification [24] to include a new species *Micromonas commoda*): a uniflagellated 2 μm organism that can be found in almost any coastal and marine environment from Arctic waters to Norweigian fjords, the coast Plymouth and even the Caribbean [25–31]. This globally dominant organism was the first reported case or viral destruction of a marine phytoplankton [32], and has established itself as a model system for host-virus dynamics since the discovery of its own dedicated lytic virus named the *“Micromonas pusilla* virus” (MPV) [33–36]. In addition *Micromonas sp*. was also the first reported case of a doublestranded RNA virus infecting a photosynthetic protist[37]. It has also been shown that lysis of *Micromonas sp*. due to the MPV increases the local DOC faster (2.5×) and larger (4.5 ×) than photosynthetic extracellular release [38]. Aside from the nutrient cycling undertaken by a cell death, the lysis of an infected cell spreads the viral infection further throughout the local population. Due to this it is reasonable to assume that organisms may have developed an avoidance response to such a hazardous circumstance yet to-date there has been minimal work on understanding how cell death could affect the motility of neighbouring cells.

Here we present a previously-unseen collective avoidance response of *Micromonas commoda* where an avoidance response is triggered upon the sudden death of a nearby cell. We first present a description of this novel response and an explanation of the underlying physical processes responsible for its activation. Next we characterise the response and compare to an analytical model of a spreading chemical patch. Finally, we reproduce the observed response by manually bursting selected cells and triggering the avoidance response in neighbouring cells.

## RESULTS

### Burst event description

Fig 1a-d outlines the cell response, which we will refer to as a “burst event”. Here we show a false-colour correlation map where any cells moving between frames will appear bright, accentuating any swimming motion in the field of view (FOV). At time *t* = 0s there are a number of moving cells in the FOV (but these are in the minority), but in the centre a single cell (Fig 1e, red) dies and/or ruptures. This releases a concentrated chemical patch into the immediate environment which proceeds to diffuse radially outwards from the cell (Fig 1f). Provided the local concentration is above the detection threshold nearby cells will detect this patch as it diffuses past them (yellow cells), triggering an avoidance response in the cells. These cells begin to swim rapidly which, when including all of the activated cells within some radius of the initial trigger cell, manifests the avoidance response as a tightly packed area of motion with a clearly defined boundary (Fig 1b). As this patch diffuses further more cells are “activated” and the event boundary continues to expand (Fig 1cg). After a longer period of time the local concentration at the edge of the patch will eventually drop below the cell detection threshold, preventing further cells from being activated and the currently active cells to escape the affected area (Fig 1d). This diffusive mechanism is demonstrated in Fig 1h with an initial Gaussian distribution diffusing over time and the detection threshold (magenta, Fig 1h insert) determining the radius of the detected patch at a time *t*. This proposed mechanism can be modelled by considering the spherically symmetric diffusion equation and an initial mass distribution.

**FIG. 1.**
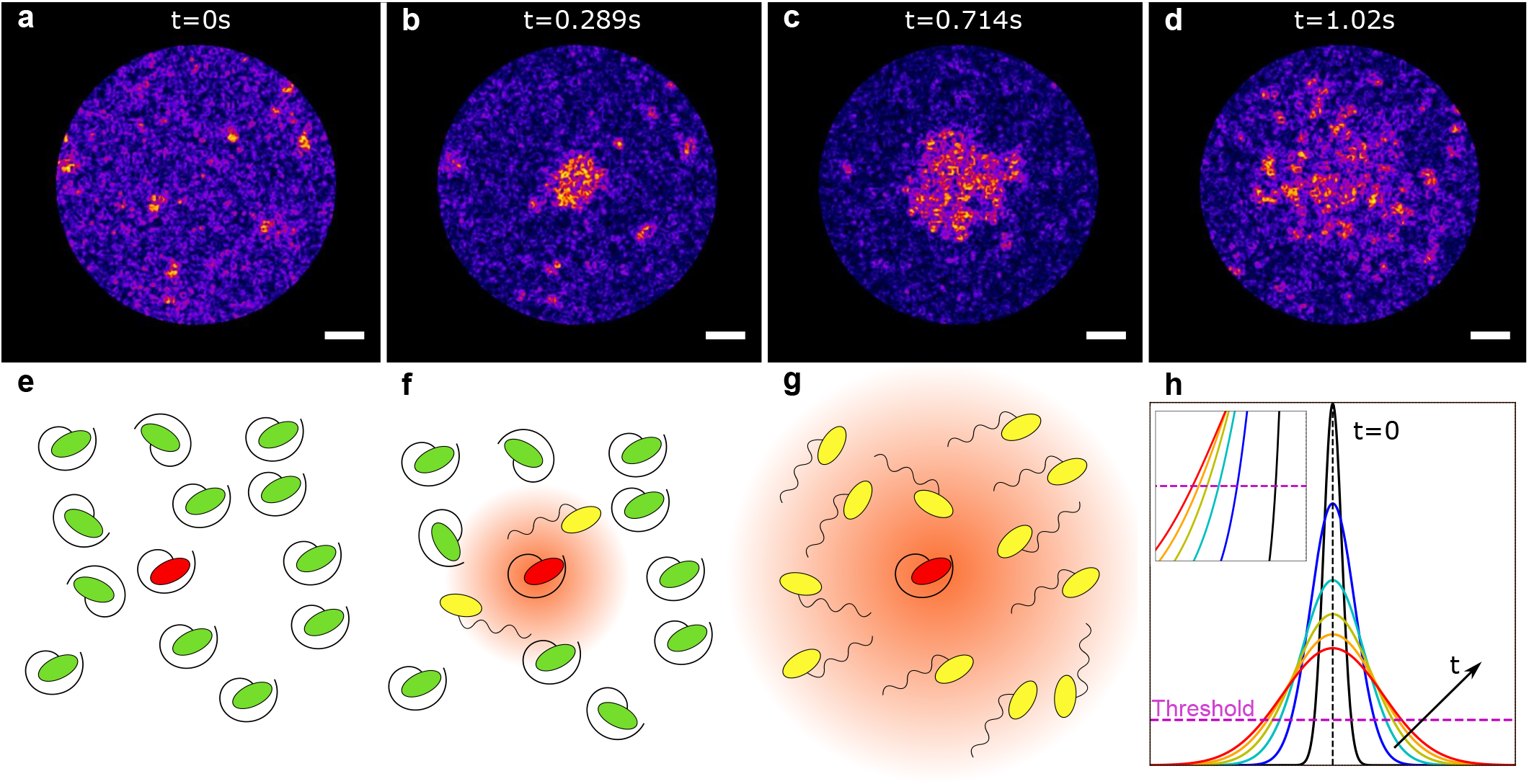
Description of a burst event. **a-d)**. False colour correlation map where motion between frames appears bright on the dark purple background. Scale bar indicates 20 *μ*m. At *t* = 0 (i.e. the instant of the event starting) there is no obvious movement aside from background cells moving. As time progresses (b,c) there is an expanding front of motion radiating out from a central point. After 1 s (d) the active cells begin to dissipate from the affected area. **e)**. Proposed event description. A trigger *M. commoda* cell (red) dies/bursts among a population of other cells (green). **f)**. This cell releases a chemical patch that is detected by the immediate neighbours (yellow) who start to swim in a phobic response to this chemical. **g)**. The patch diffuses further, activating more cells in the local area. **h)**. Diffusion from an initial Gaussian distribution, where the detection threshold of the cells will dictate the detected radius of the spreading patch (insert).

### Spherical diffusion model

Assuming a radially symmetric initial distribution and uniform diffusion in all directions we can reduce to the spherically symmetric diffusion equation:

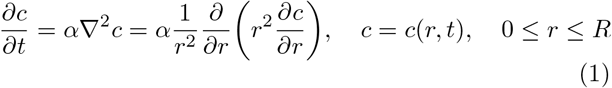

Where the concentration of the chemical stimulant at a distance *r* from the initial source at a time *t* is given by *c*(*r, t*). The initial boundary conditions are defined as:

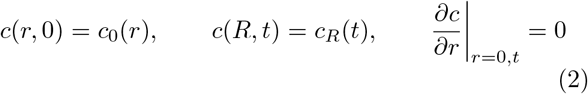

This can be solved by separation of variables to obtain an expression describing how the mass of the chemical will diffuse out from the initial mass profile *c*_0_(*r*). For simplicity, we define this initial profile to be a Gaussian profile with a full width half maximum (FWHM) of 1.5 × 10^-6^ m with a total mass M, approximately the size of *Micromonas sp*. Further assuming an instantaneous mass release and setting the background concentration *c_R_*(*t*) = 0 (since shifting this background can be thought of as effectively shifting the detection threshold of the cell), we define the initial conditions:

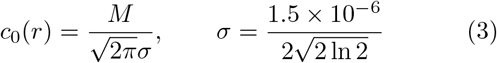

There are three main quantities that will determine the spreading dynamics of the chemical: the mass released (*M*), the diffusion coefficient of that chemical (*D*) and the detection threshold of the organism (Γ). Unfortunately we do not know the chemical stimulus that is responsible for triggering these events, so for the purpose of this validation model we instead make some biologically relevant estimates for the above quantities using known values for dimethlysulfiopropionate (DMSP). DMSP is a solute released by plankton as a specific point source during events such as cell lysis and grazing and in cells similar to *Micromonas sp*. (in phytoplankton populations) contain 45.5±73.7 fg of this solute. Assuming the average value, and that all of this mass is released instantly, we set *M* = 4.5 × 10^-17^ kg. The detection threshold (Γ) is set at 10^-6^ M which is in line with the DMSP detection threshold for bacteria who operate at the same length scale as *Micromonas sp*. The diffusion coefficient (*D*) is left as a free-fitting parameter, though we expect a value within the range of 10^-11^ — 10^-9^ m^2^s^-1^ based on the diffusion coefficient of other particulates. Any enhancement effects of nearby cell motion on the diffusion coefficient is not treated separately since there is no information on the size/shape of the pollutant particles. Finally, we define the patch radius *r_p_*(*t*) to be the radius at a time *t* that contains 95% of the initial mass *M*.

### Experimental radius and patch radius

Fitting our experimental model patch radius *r_p_*(*t*) to the experimentally measured radius *r*(*t*) produces the red curve shown in Fig 2, with *D* = 1.14 m^2^s^-1^. For the first second of the event the model radius faithfully reproduces the observed behaviour, but diverges after this period. However this is to be expected since (as shown in Fig 4d-g) the event tracking begins to breakdown after 1 s as there is no longer a coherent front to track. Indeed the radius measurements here will jump between the remnants of the patch and the free swimming cells which have a free-swimming speed [39] of *v* ≈ 23 *μ*m.s^-1^ as labelled on Fig 2. Here we see that after 1 s the event radius *r*(*t*) takes a value between the patch radius and the radius dictated by the swimming speed of the cells.

**FIG. 2.**
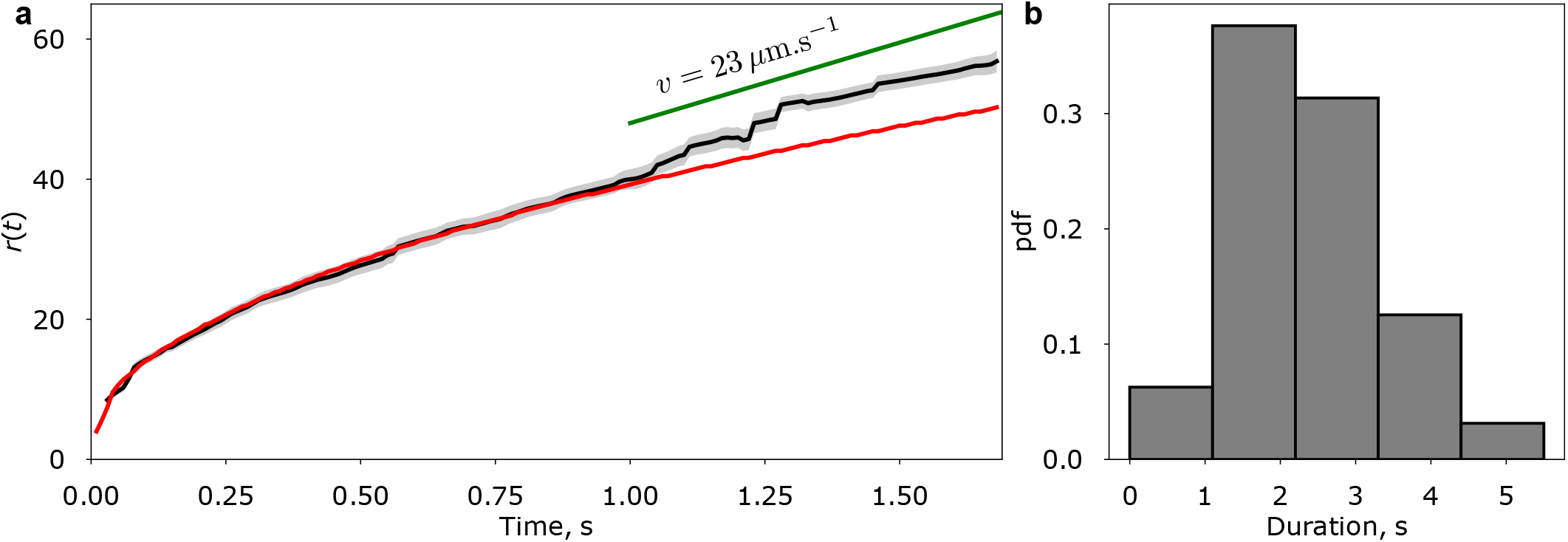
Burst event characterisation. **a)**. (Black) The experimentally measured average radius *r*(*t*) from 20 individual events with a shaded standard error. (Red) The model patch radius *r_p_*(*t*) with the values: *M* = 4.6 × 10^-17^ kg, Γ = 10^-6^ M with a fitted diffusion coefficient *D* = 1.1 × 10^-11^ m^2^s^-1^. For the first one second of the events the patch model faithfully reproduces the observed behaviour, then diverges as the swimming speed of the cells on the edge of the event front exceeds the diffusive speed of the patch. **b)**. Distribution of the event duration for 50 events, which is defined as the time taken for the activity in the affected region to return to background levels (see Fig 4c).

A point should be made here about the validity of this model. The resultant curve produced from the model patch radius can be achieved by fixing any two of *D*, Γ, *M* and leaving the third as a free fitting parameter. It should be stressed that this model is being used as a means of verifying that the detection of a realistically sized pollutant patch diffusing from a local source can produce the same radius expansion behaviour as is observed in the experiments, and is not being used to make predictions on these physical quantities.

### Laser ablation to trigger response

To this point, all of the recorded events have been events that occurred spontaneously in the millifluidic device in the absence of any form of external stimulus. To confirm that the death/bursting of a cell is responsible for triggering the avoidance response of nearby cells we targeted specific cells with a laser ablation apparatus. Fig 3 displays the trajectories from two experiments: (a-d) a control where the laser was targeted on a point adjacent to a cell and (e-f) where a cell was directly targeted. The image processing used to produce these images are described in more detail in Methods but in short for a frame at time *t*_1_, a bright spot indicated a cell has moved through that spot in the interval *t* ∈ [0,*t*_1_], hence particle trajectories will appear as bright tracks. In the control experiment (Fig 3a-d), the laser targets a point (red cross) adjacent to a cell (cyan circle). In the subsequent frames, there is no obvious response from surrounding cells bar diffusive motion. In the second experiment (Fig 3e-h) a single cell was directly targeted by the laser. Immediately after ablation there is clear motion in the surrounding cells in comparison to the control, confirming that the rupturing of a single *Micromonas sp*. cell is sufficient to trigger an avoidance response in the surrounding cells.

**FIG. 3.**
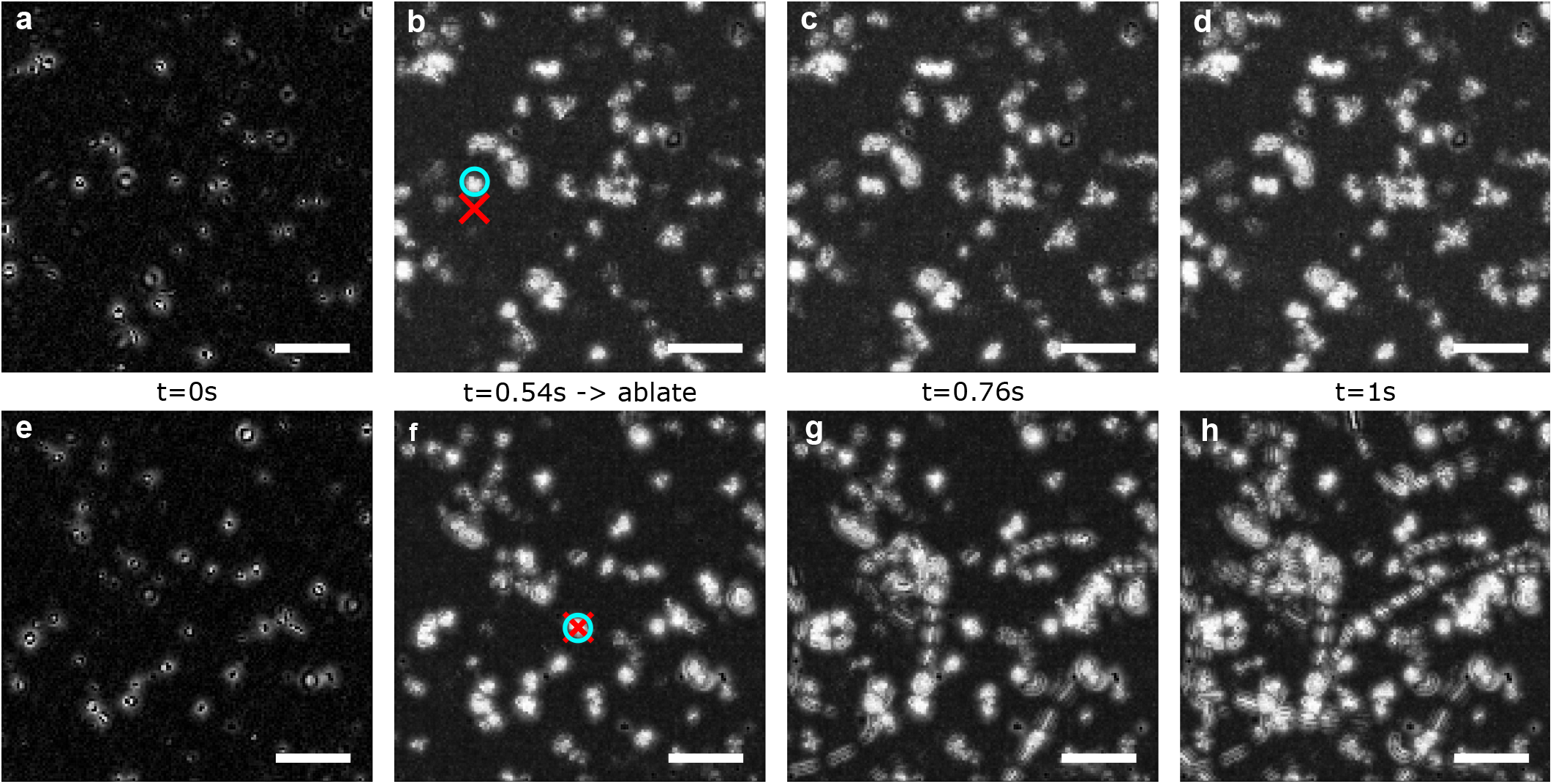
Manually event trigger using laser ablation. Bright areas indicate that a cell has moved through that location during the last *t* seconds. The ablation takes place at *t* = 0.54 s and targets the red cross. The selected target cell is circled in cyan. Scale bars indicate 20*μ*m. **a-d)**. Control experiment, where the laser targets (red) space adjacent to the cell (circled blue). There is no obvious signs of motion after the pulse. **e-h)**. The laser now directly targets a cell (in a new region), and in the subsequent frames nearby cells exhibit the previously seen avoidance response.

## DISCUSSION

The death of phytoplankton organisms in the oceans has a profound impact on both the local conditions and the global ecosystem. These organisms are responsible for half of the global oxygen production during their life cycle and release large amounts of DOM into the environment when the cell dies. The oceans are estimated to contributing to half the primary productivity of the entire biosphere, and since living phytoplankton form approximately quarter of the carbon mass in the oceans the local chemical changes induced by a single cell death are not to be underestimated. On a small scale cell death provides a food source for nearby organisms but can also act to alert nearby cells to danger sources through the release of chemical signals e.g. DAMPs. One of the most populous members of the phytoplankton community is the pico-eukaryote *Micromonas sp*., a uni-flagellated single celled organism that has been reported in a wide range of marine and coastal environments across the globe. This organism has become a model system for studies of host-virus dynamics and viral lysis, though previous studies have not investigated the effect cell death has on the motility on nearby cells.

Here we have presented a novel avoidance response where the death of a single cell of *Micromonas sp*. triggers an avoidance response in the surrounding population. During the death of the organism a chemical is released that diffuses radially out from the cell, which when detected in sufficient concentration triggers nearby cells to immediately begin swimming rapidly to escape the local environment and any hazard it contains. On a larger scale this response manifests itself as a “burst event” where an event front of cells being activated/triggered propagates radially outwards from the source cell. The duration of these events were measured by analysing the PIV velocity in concentric rings centred on the origin of the event, and the evolution of the event radius measured by developing a image processing workflow combined with a circular feature recognition algorithm. Modelling a diffusing patch with biologically relevant quantities verifies that a single *Micromonas sp*. cell is capable of exuding a chemical patch sufficient to reproduce the observed event behaviour. Finally, manually ablating a cell is shown to also trigger the same individual cell behaviour underlying the burst event dynamics.

This novel response presents a number of avenues for future exploration starting with the frequency of these events and the nature of the chemical source. During the experiments recording the events it was noted that the events occurred at an increasing rate the longer the cells were contained in the device. It was also seen that renewing the media surrounding the cells before filling the device led to a reduction in the rate, but more significantly was that chambers that were kept in dark conditions for several hours prior to filming had a dramatic fall in the event rate compared to samples that were illuminated. These preliminary results (not shown here) suggest that buildup in metabolic waste in the sample could affect either (or both) influence the sensitivity of cells or increase the ambient concentration of the responsible chemical signal as to increase the event rate when the cells are more active i.e. during the day phase. Lastly, further work should also be done to attempt to isolate the chemical signal responsible and compare with known DAMPs to determine if the cell is releasing a specifically produced compound to alert nearby cells or if the organism has evolved to interpret the sudden release of DOM from the source cell as a trigger itself.

## METHODS

### Cell culturing and experiments

Cultures of *Micromonas commoda* (RCC827) were maintained in Guillard’s f/2 medium [40], prepared using artificial seawater (Sigma) and f/2 nutrient solution (Sigma) in 500 ml quantities excluding the sodium-glycerophosephate to reduce any precipitation in the medium. The cultures were grown in a diurnal chamber on a 16/8 hour light/dark cycle at a constant temperature of 20°C. A Nikon TE-2000U inverted microscope with a longbandpass filter (765 nm cutoff wavelength, Knight Optical UK) was used to limit any phototactic behaviour due to the imaging light. Cells were filmed (Allied Vision Pike camera) at a variety of magnifications but typically at 20 × to maintain single particle recognition with a wide field of view (370 μm). Cells were loaded into a cuboid polydimethysiloxane (PDMS) millifluidic chamber with internal dimensions 10 mm × 10 mm × 4 mm plasma bonded to a class cover slip (Harrick) and left to sediment in the diurnal chamber for several hours before filming commenced.

Events were captured by filming as continuously as possible for several hours to capture as many spontaneous events as possible, typically at 30 FPS. The duration of events was calculated from a larger data set including events recorded with reduced framerates (10FPS) and included events that were not sufficiently optically resolved to track reliably but were still of enough quality to determine the start/end points respectively.

### Laser ablation

Images were collected in brightfield mode on an inverted spinning disk confocal microscope (Marianas SDC, 3i) using a40× objective (1.3 NA, Zeiss) at 50FPS. Ablation was performed by targeting individual algae with a single 1.3 ns pulse from a 532 nm passively Q-switched laser (Ablate, 3i).

### Image processing

The false-colour images shown here were produced using ImageJ [41] using an external plugin [42] to correlate each frame with the subsequent frame. After this the contrast was enhanced and normalised (using default tools) and the feature edges located (using default tools), the process of which was kept consistent across all frames. The same process was utilised for the images in Fig 3 apart from a background subtraction tool was used after the original processing which was implemented consistently across all processed frames. These images have reduced contrast compared with the other false-colour images due to the reduced cell density in the ablation experiments, hence are shown in grayscale to (1) improve the image quality and to (2) signify the additional image processing.

### Locating the origin of the event and calculating the event duration

Locating the event origin (both spatial and temporal) is a surprisingly involved affair – if we ignore the simple “by eye” approach. A method was developed that took advantage of the fast nature of these events. Each brightfield image was divided into a square grid (typically 10 × 10), then each square was correlated with itself but separated by a number of timesteps dependent on the individual experiment framerate. We follow the approach of Dikbas et al. [43] to define two correlation coefficients based on the horizontal and vertical correlation respectively. A two-dimensional approach was used to maintain analytical robustness in the cases where the local cell distributions were distinctly inhomogenous (since the event tracking inherently relies on the local spatial distribution of cells). For a fuller description please refer to the source material, but in short we define horizontal (vertical) coefficients *C_h_* (*C_v_*) for two *m* × *n* matrices **A, B**:

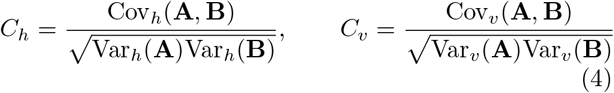

Where the variance/covariance (Var, Cov resp.) are defined as:

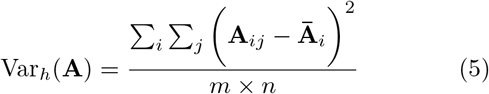

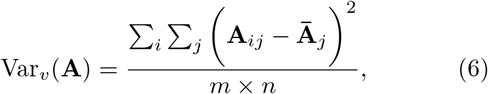

and

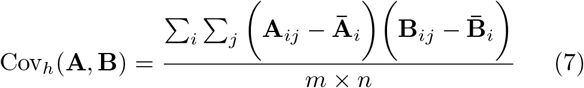

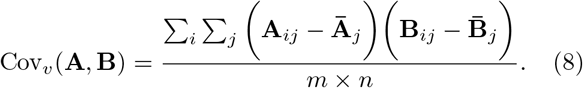

Here **A** is the grid square at a time *t* and **B** the same grid square at a time *t* + Δ*t*. For a expanding circular event, *C_h_* = *C_v_* so comparing the two can give some measure of any asymmetry of the event. Due to the circular nature of the events, the first grid square to drop in correlation value should contain the origin of the event, as shown in Fig 4b. Identifying the time of this drop will identify the start time of the event (Fig 4b inset) and the spatial origin can be determined by repeating this correlation method with increasingly small grid squares until a square is found such that the size of the square matches the diameter of the event after one timestep. The centre of this final square is taken as the spatial origin of the event.

**FIG. 4.**
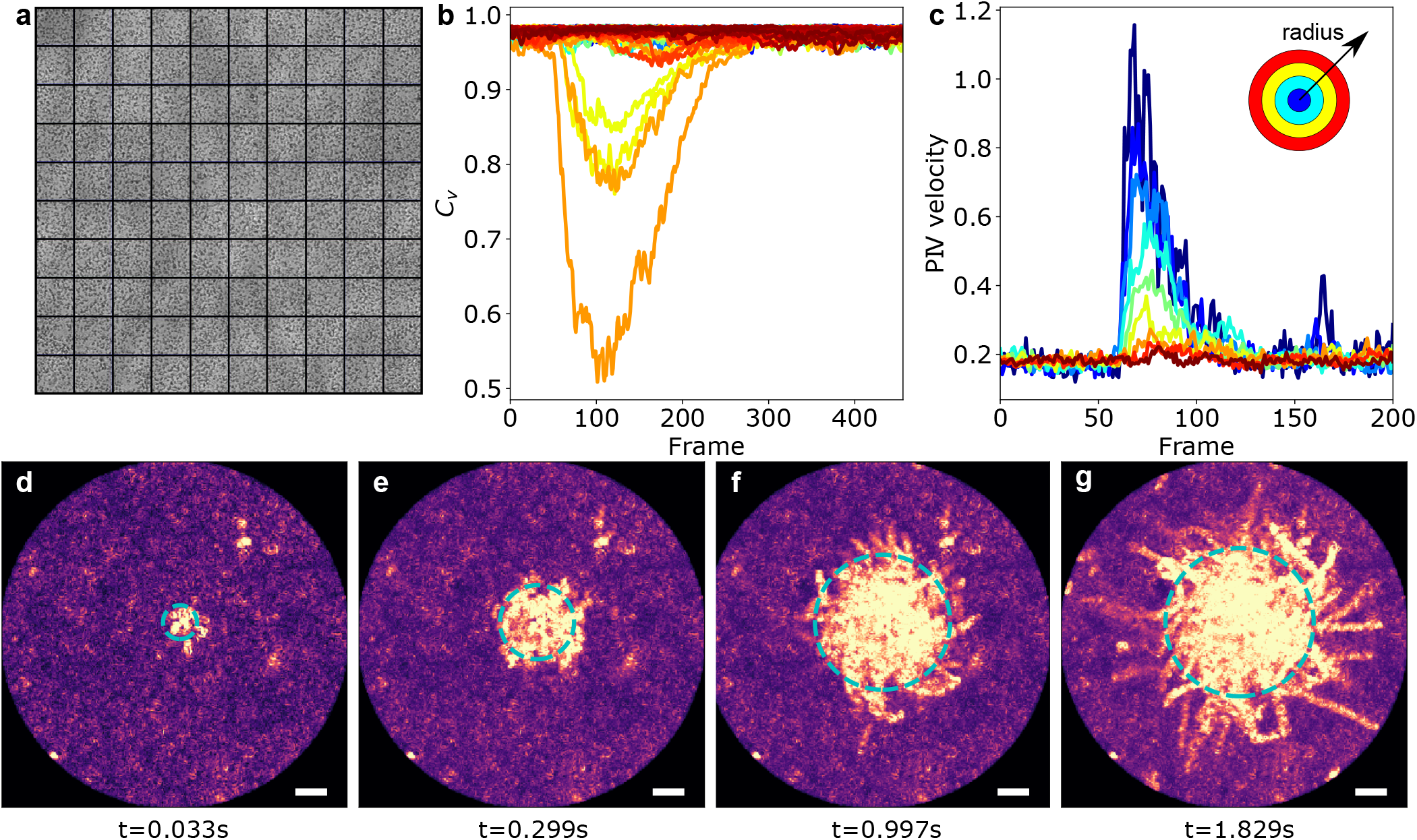
Methods for characterising the events. **a)**. The initial brightfield image, where cells appear as dark spots. This image is divided into a 10 × 10 grid and each grid square is correlated with itself at time *t* and *t* + Δ*t*. **b)**. Vertical correlation (*C_v_*) plot for each grid square as a function of time (measured in frames). The affected squares are immediately evident, and comparison of the start times of each square drop (insert) identifies the square containing the spatial origin. This process is repeated with smaller grids until the origin is identified to the nearest pixel. **c)**. Analysing the PIV velocity radially from the origin enables calculation of the event duration from the start of the spike in velocity to when the velocity in the affected area returns to background values. **d-g)**. Plots of **V**_*ijk*_ for four different times. Bright areas indicate there has been a moving cell in that spot at some point in that spots history. As time progresses, the event front boundary is clear and easily detectable by implementation of a circular feature recognition algorithm until (g) when the swimming speed of the cells activated at the boundary exceeds the diffusive speed of the patch and/or the local chemical concentration drops below the cell’s detection threshold. Scale bars indicate 20 *μ*m.

The event duration is defined as the time taken from the start of the event for the cell activity in the local region of the event to return to normal. This can be characterized by performing particle image velocimetry (PIV) on the affected area, then examining the resultant velocity in concentric rings centered on the origin of the event (Fig 4c. From this we calculate the end of the event to be when the PIV velocity in all the affected rings returns to a comparable mean and standard deviation before the event started.

### Event front tracking

Examining the distribution of active cells in Fig 1, we see on timescales around 1 s the burst event has a reasonably well defined edge between activated cells and nonactivated cells. This edge, which when measured from the centre of the event, defines the event radius *r*(*t*), and can be measured with the combination of an image maximization technique and a circular feature recognition algorithm. We define the matrix **T**_*ij*_ (*t*) as the two-dimensional matrix representing the intensity of the image at location (*i,j*) at a time *t*. Then we define the following three-dimensional matrix:

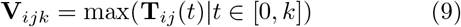

Or explicitly for each timestep, each element of **V** are equal to the maximum value it attains in any of the previous timesteps. This produces Fig 4d-g where we see the centre of the event now remains bright as opposed to fading as the event progresses. An interesting side effect of this method is that individual tracks of cells remain bright and so potentially offers a less computationally expensive method of PTV analysis. A circular feature recognition algorithm (MATLAB) can now reliably detect the edge of the event and so measure the event radius *r*(*t*).

## ACKNOWLEDGEMENTS

The authors gratefully acknowledge Dr. Joseph Christie-Oleza for kindly supplying the initial cultures of *Micromonas commoda* and Dr. Claire Mitchell of CAMDU (Computing and Advanced Microscopy Unit) for their support and assistance in this work. RH acknowledges EPSRC Award 1619257 for funding.

## AUTHOR CONTRIBUTIONS STATEMENT

RH and MP devised the study; RH and JR carried out experiments; RH undertook data analysis and presentation of results; RH and MP wrote this manuscript.

## ADDITIONAL INFORMATION

The author(s) declare no conflicting interests.

